# Auditory High Entropy Response (A-HER): A Novel Paradigm for Studying Brain Processing of Uncertain Information

**DOI:** 10.1101/2023.12.27.573480

**Authors:** Xiaoqi Liang, Qianyun Zhu, Zhiguo Zhang, Zhenxing Hu, Zhen Liang, Linling Li, Li Zhang, Xuezhen Xiao, Gan Huang

## Abstract

This paper introduces a novel experimental paradigm - Auditory High Entropy Response (A-HER), which maximizes the information entropy of auditory stimulus sequences. This allows us to study how the brain processes complex information, rather than isolated individual events. Our analysis of the frequency response of the frontal theta rhythm induced by A-HER indicated a significant increase in signal-to-noise ratio and repeatability compared to zero-entropy Auditory Steady-State Response (A-SSR) and low-entropy mismatch negativity (MMN). We further investigated whether the A-HER response was induced by stimulus sequence differences or uncertainty, and studied its propagation rules. Different principles between evoked and entrained were found in A-HER and A-SSR. In conclusion, the A-HER paradigm, by maximizing stimulus sequence uncertainty, offers a new approach to analyzing how the brain processes uncertain information. It has potential for diagnosing and researching neurological and mental diseases, and for brain-computer interfaces, thus potentially impacting neuroscience, cognitive science, and psychology.

## Introduction

The quest to unravel how the human brain grapples with uncertain information stands as a cornerstone in the field of cognitive neuroscience studies (***Hsu et al., 2005; White et al., 2019; Summerfield, 2022; Soltani and Izquierdo, 2019; Mulders et al., 2023***). Conventional experimental paradigms have historically centered around scrutinizing isolated events or solely stimulation sequences, thereby limiting our comprehension of the brain’s intricate operations amidst complexity and uncertainty. For instance, in auditory experiment, Auditory Evoked Potentials (AEPs) (***Burkard et al., 2007***) isolate and examine single stimuli, restricting the exploration of higher-level cognitive processes linked to uncertainty. While Mismatch Negativity (MMN) (***Garrido et al., 2009***) detects deviations in auditory sequences, its capacity to unravel the brain’s comprehensive response to uncertainty within diverse sequences remains constrained. The usefulness of the Auditory Steady-State Response (A-SSR) (***Spencer et al., 2008; Korczak et al., 2012***) in studying auditory entrainment is notable, yet its limitations are evident in capturing the brain’s dynamic intricacies when facing varying uncertainty levels within sequences. Thus, broadening experimental methodologies beyond isolated events and solely stimulation sequences is crucial for a more comprehensive understanding of how the human brain navigates uncertain information scenarios.

In neuroengineering, the Steady-State Response (SSR) (***Herrmann, 2001; Spencer et al., 2008***) represents a significant event-related potential (ERP) evoked by continuous sensory stimuli, widely employed for assessing sensory function and diagnosing processing dysfunctions (***Vialatte et al***., ***2010; Thuné et al., 2016; Korczak et al., 2012***). Notably, the Visual Steady-State Response (V-SSR), or Steady-State Visual Evoked Potential (SSVEP), stands out in Brain-Computer Interface (BCI) applications due to its impressive Signal-to-Noise Ratio (SNR) and high Information Transfer Rate (ITR) (***Chen et al., 2015; Nakanishi et al., 2017***). However, the fatigue induced by SSVEP-based BCIs can lead to user discomfort, signal quality deterioration, system performance degradation, and potentially increase the risk of photosensitive epileptic seizures, thus significantly constraining their use(***Vialatte et al., 2010***). Therefore, exploring non-visual paradigms within BCI research is imperative. Despite demonstrations of selective attention effects in auditory information processing, the limited magnitude and extended trial time (> 40 sec) of A-SSR hinder its viability in BCI (***Kim et al., 2011; Lopez et al., 2009***). Enhancing the SNR in auditory stimuli responses remains a critical objective for advancing auditory-driven BCIs (***Akcakaya et al., 2013***).

Emerging from the quest for a more comprehensive understanding of uncertain information processing in brain, the Auditory High Entropy Response (A-HER) paradigm is proposed as a pioneering approach in this work. Unlike previous methods, which focused on isolated events or solely stimulus sequences, A-HER deliberately maximizes information entropy within auditory stimuli. This deliberate emphasis offers a promising avenue to delve into the brain’s handling of intricate, uncertain, and information-rich auditory scenarios. Furthermore, the newly proposed A-HER paradigm shows potential in averting Repetition Suppression (RS) (***Todorovic et al., 2011***) effects and enhancing the SNR, thereby amplifying its relevance in neuroengineering applications.

### Auditory High Entropy Response (A-HER)

Entropy is a scientific concept as well as a measurable physical property that is commonly associated with a state of disorder, or randomness (***Clausius, 1865; Dugdale, 2018***). In information theory, Shannon entropy (***Lin, 1991***) is a measurement of the uncertainty of a sequence. Considering two types of stimuli in a sequence, like the standard (***S***) and deviant (***D***) stimuli for oddball paradigm, the entropy can be calculated as

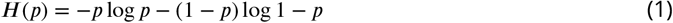

in which *p* is the prior probability for stimulus ***S***) in the sequence. Based on the definition of entropy in Eq.(1), we can calculate the entropy for different types of sequences, as illustrated in Fig. 1.

**Figure 1.**
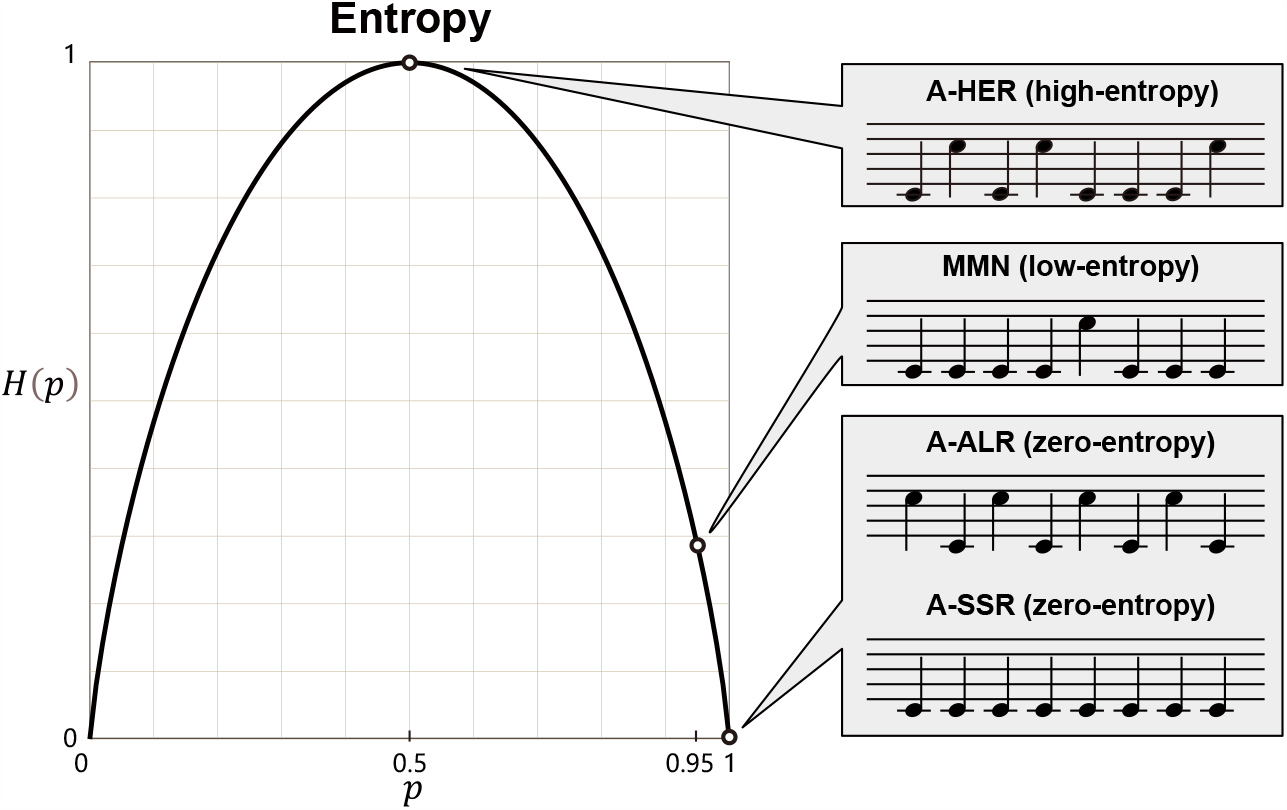
The auditory stimulus sequences in A-HER, MMN, A-ALR, and A-SSR paradigm with their entropies. For A-SSR, with the prior probability p=1 for the standard stimulus, we have the entropy of the whole sequence *H*(*X*) = 0. For A-ALR, with two types of stimuli appearing alternatively, the whole sequence is also deterministic with the entropy *H*(*X*) = 0. For MMN, the standard stimuli occur with a larger probability *p* = 0.95, so the entropy of the whole sequence is also small *H*(*X*) = 0.29. For A-HER, two types of stimuli occur with the equal probability *p* = 0.5, so the entropy reaches the maximum *H*(*X*) = 1.

- **A-SSR**: With the prior probability of the standard stimulus *p* = 1, the entropy *H*(*p*) = 0 indicates that A-SSR is evoked in a fully deterministic sequence with zero-entropy.
- **A-ALR**: Auditory Alternative Response (A-ALR) is another deterministic sequence. Both two types of stimuli still termed ***S*** and ***D***, come alternately with the prior probability *p* = 0, or 1. Hence, A-ALR is evoked in a zero-entropy sequence with *H*(*p*) = 0.
- **MMN**: With a random uneven probability appeared with the stimulus ***S*** (*p* = 0.95) and ***D*** (1−*p* = 0.05), for example. MMN is evoked in a low-entropy sequence with *H*(*p*) = 0.29.
- **A-HER**: If the stimuli ***S*** and ***D*** appear randomly with the same probability *p* = 0.5, the stimulus sequence with the highest entropy *H*(*p*) = 1, achieves the largest uncertainty.

## Experiment design

A total of 23 healthy subjects (13 women; mean age 24.9 years, ranging from 22 to 39 years) participated in the three experiments. All participants are non-musicians and right-handed. And all reported normal hearing, normal or corrected to normal vision, and no history of neurological or psychiatric disease (as indicated in a self-report).

Before the experiment, all subjects were informed of the experimental procedure and signed informed consent documents. Ethical approval of the study was obtained from the Medical Ethics Committee of the Health Science Center, Shenzhen University (No. PN_2021-035).

For the newly proposed A-HER paradigm, three experiments were designed to answer the following three questions:

1. Can high-entropy stimulus sequences in A-HER alleviate the repetition suppression effect and bring a higher response than A-SSR? If so, what are the characteristics of its frequency response and the distribution of its topographic map?
2. Within the A-HER paradigm, the crucial factor driving maximal response is the discrepancy among stimuli or the inherent uncertainty?
3. Comparing the induction methods of A-HER and A-SSR, what would be the differences in their induction principles?

The details of the three experiment paradigms were illustrated in Fig. 2.

**Figure 2.**
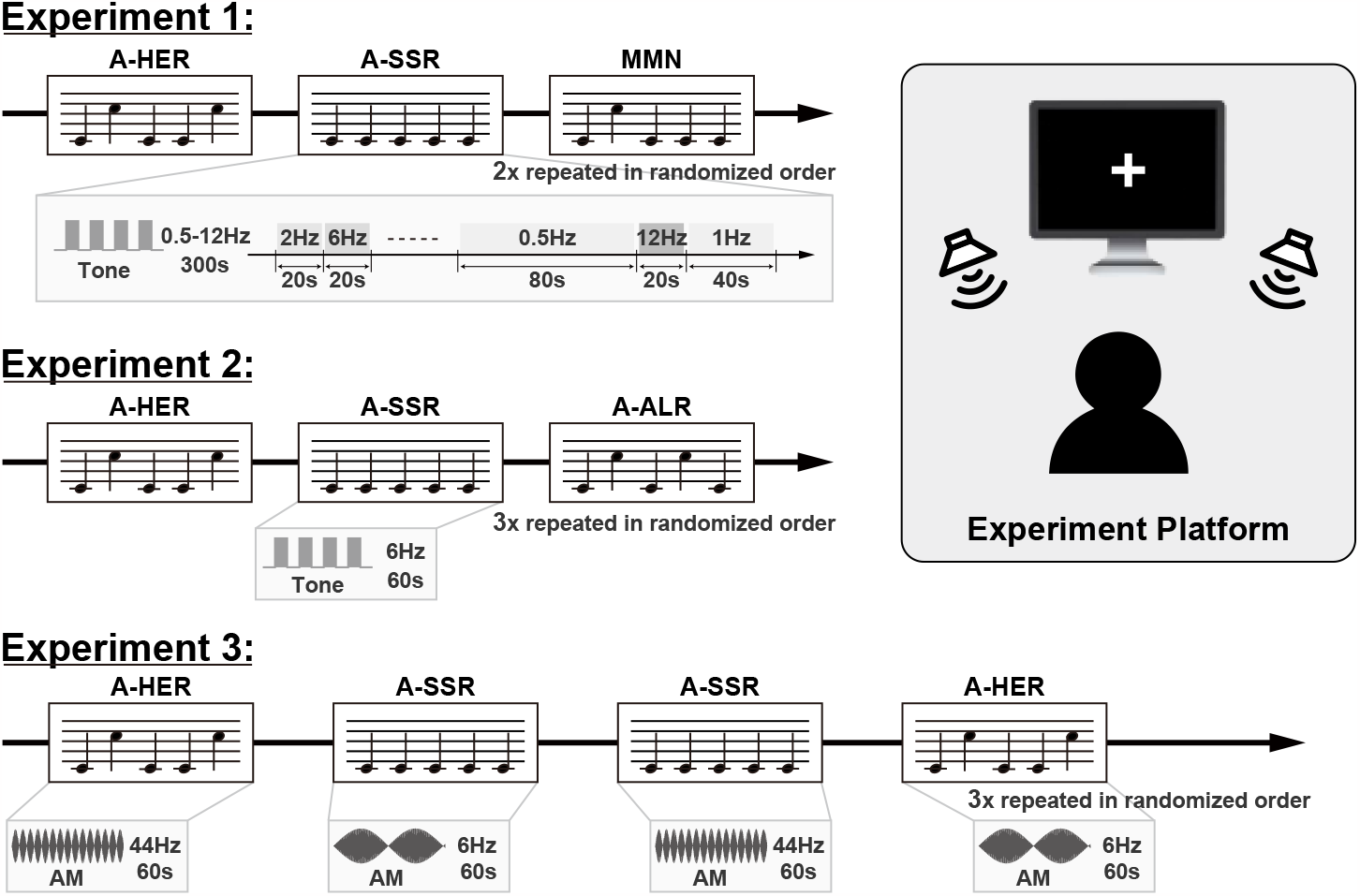
The auditory stimulus sequences in A-HER, MMN, A-ALR, and A-SSR paradigm with their entropies. For A-SSR, with the prior probability p=1 for the standard stimulus ***S***, we have the entropy of the whole sequence *H*(*X*) = 0. For A-ALR, with two types of stimuli appearing alternatively, the whole sequence is also deterministic with the entropy *H*(*X*) = 0. For MMN, the standard stimuli ***S*** occur with a larger probability *p* = 0.95, so the entropy of the whole sequence is also small *H*(*X*) = 0.29. For A-HER, two types of stimuli occur with the equal probability *p* = 0.5, so the entropy reaches the maximum *H*(*X*) = 1.

### Experiment 1

With different stimulation frequencies, the temporal, spatial, and frequency distribution characteristics of A-HER are investigated as compared with A-SSR, and MMN. For each block of A-SSR, MMN and A-HER, 11 types of stimulation frequencies, including 0.5, 1, 2, 3, 4, 5, 6, 7, 8, 10, and 12 Hz, are applied in random order. The stimulation durations corresponding to the stimulation frequencies are 80, 40, 20, 20, 20, 20, 20, 20, 20, 20, and 20 seconds, respectively. For low-frequency (0.5 Hz and 1 Hz) stimulation, the duration of the stimulation is set to be longer to ensure that there were at least 20 stimuli for each stimulation frequency in a block. Hence, the total stimulation duration of each block is 300 seconds. The blocks of A-SSR, MMN, and A-HER are arranged in random order and repeated two times. The subjects could have a rest as they wish between two blocks.

### Experiment 2

A-HER was compared with A-SSR and A-ALR to examine whether A-HER is evoked by the difference or uncertainty among the stimuli. The stimulus frequency was fixed at 6 Hz. The blocks of A-SSR, A-ALR and A-HER were delivered in a 60-second period. The three types of blocks were arranged in random order and repeated three times.

### Experiment 3

Auditory stimulation of pure tone bursts (Tone) in Experiments 1 and 2 were compared with amplitude modulated (AM) stimulation patterns. For the AM-based auditory stimulation, carrier frequency was set as 524 Hz or 262 Hz for standard (***S***) or deviant (***D***) stimuli, modulation frequency was set as 6 Hz or 44 Hz in different blocks of A-SSR and A-HER. Similarly, the blocks of A-SSR and A-HER were delivered within during of 60 seconds. The four types of blocks were arranged in random order and repeated three times.

## Results

### Experiment 1

To investigate the frequency response features of the newly proposed A-HER paradigm, the EEG responses of A-HER, A-SSR, and MMN in Experiment 1 were compared in both the time domain and frequency domain (Fig. 3).

**Figure 3.**
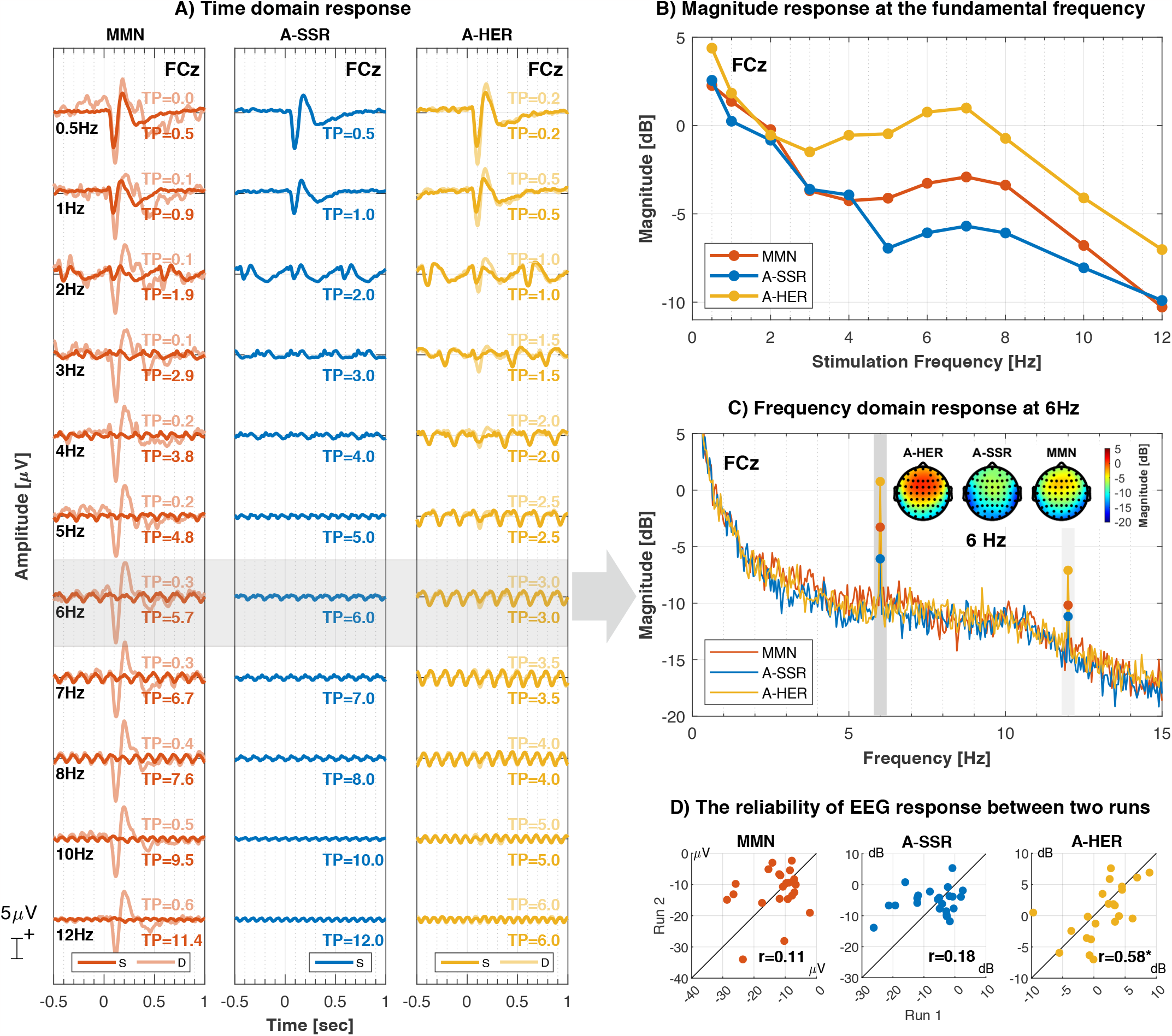
The grand averaged time domain response of A-HER, A-SSR, and MMN in Experiment 1. (A) The ERP response of A-HER(S), A-HER(D), A-SSR(S), MMN(S), and MMN(D) with different stimulation frequencies from 0.5 to 12 Hz. “TP” indicates the temporal probability, which is the average number of times for one type of stimulus occurring in one second. “S” and “D indicates standard and deviant stimuli. (B) The peak-to-peak magnitude of these five types of ERP with different stimulation frequencies (C) The detail of the time domain response of the five types of ERP response at the stimulation frequencies 6 Hz, with their topographies of their peaks and troughs.

#### Time-domain response

The time-domain responses of standard (***S***) and deviant (***D***) stimuli, as depicted in paradigms A-HER, A-SSR, and MMN at channel FCz, are illustrated in Fig. 3A. As the stimulation frequencies fluctuate from 0.5 to 12 Hz, the stimulus onset synchrony (SOA) diminishes from 2 to 0.083 seconds.

For the standard stimuli (***S***), a paired two-sample *t*-test showed no significant difference between the peak-to-peak responses of A-HER (13.31 *μ*V) and MMN (13.51 *μ*V, *p* = 0.53), as well as A-SSR (13.22 *μ*V, *p* = 0.40) at 0.5 Hz. As the stimulation frequency escalates, all three paradigms display changes in the magnitude of the peak-to-peak response, albeit with unique trends. For instance, at a stimulation frequency of 6 Hz, the magnitude of A-HER (4.09 *μ*V) is significantly higher than that of MMN (2.43 *μ*V, *p* = 2.95 × 10^−8^) and A-SSR (1.31 *μ*V, *p* = 2.66 × 10^−9^). Similarly, at a stimulation frequency of 12 Hz, A-HER (1.62 *μ*V) significantly exceeds MMN (1.20 *μ*V, *p* = 7.53 × 10^−7^) and A-SSR (0.83 *μ*V, *p* = 7.50 × 10^−10^) in magnitude.

For the deviant stimuli (***D***), the peak-to-peak response at 0.5 Hz is 19.92 *μ*V for A-HER and 20.86 *μ*V for MMN, which are significantly larger than the magnitude for the standard stimuli (***S***) with the paired two-sample *t*-test *p*-value *p* = 5.37 × 10^−9^ and *p* = 8.28 × 10^−8^ respectively. As the stimulation frequency increases, the magnitude of the deviant stimuli (***D***) in MMN reduces to 19.84 *μ*V at 6 Hz, and further to 15.91 *μ*V at 12 Hz, which are significantly larger than the magnitude for the standard stimuli (***S***) in MMN with the *p*-value *p* = 5.45 × 10^−12^ and *p* = 6.16 × 10^−13^, respectively. For A-HER, the average magnitude decreases to 4.26 *μ*V at 6 Hz, and 1.75 *μ*V at 12 Hz for the deviant stimuli (***D***). Compared with the magnitude for the standard stimuli (***S***) in A-HER, there is no significant difference (*p* = 0.20 at 6 Hz and *p* = 0.13 at 12 Hz) in the paired two-sample *t*-tests.

Furthermore, it should be noticed that with the same temporal probability TP=0.5, the magnitude of deviant stimuli (***D***) in MMN at 10 Hz is 17.23 *μ*V, which is significantly larger than that of standard stimuli (***S***) 13.22 *μ*V in A-SSR at 0.5 Hz (*p* = 1.83 × 10^−7^).

#### Frequency-domain response

The frequency domain responses of A-HER, A-SSR, and MMN paradigms with different stimulation frequencies at channel FCz is shown in Fig. 3B. Different from ERP analysis, the frequency domain Fourier Transform is unable to discriminate between the standard (***S***) and deviant (***D***) stimuli’s responses. For low stimulation frequencies, such as 0.5 Hz, 1 Hz, and 2 Hz, the large SOAs prevent the establishment of a stationary modulation of the EEG rhythm. The spectra of the responses observed in the frequency domain are predominantly characterized by the frequency characteristics of their ERPs, which concentrate mainly in the delta and theta frequency bands. As illustrated in Fig. 3B, the magnitudes at their respective fundamental frequencies appear similar. As stimulation frequency increases, A-HER demonstrates a higher response than A-SSR and MMN. All three paradigms reach a local maximum at theta band, a result that aligns with the time domain response in Fig. 3A.

Fig. 3C presents the frequency domain response of A-HER, A-SSR, and MMN with a stimulation frequency of 6 Hz. The paired-sample *t*-test result reveals that the magnitude of A-HER is significantly higher than that of A-SSR (*p* = 1.46 × 10^−5^) and MMN (*p* = 3.16 × 10^−4^) at the fundamental frequency of 6 Hz. At the harmonic frequency of 12 Hz, the magnitude of A-HER is also significantly greater than A-SSR (*p* = 2.85 × 10^−4^) and MMN (*p* = 1.21 × 10^−3^). All three paradigms display similar topographies centered around FCz, albeit with different magnitudes.

#### Reliability analysis

The reliability test was conducted on the EEG responses from two separate runs using Spearman correlation analysis for all three paradigms depicted in Fig. 3D. For the MMN paradigm, the difference between the responses to the deviant (***D***) and standard (***S***) stimuli at 0.5 Hz was found to have a correlation coefficient of r=0.11 (*p* = 0.61). For the A-SSR and A-HER paradigms, the frequency responses at their fundamental frequency of 6 Hz revealed correlation coefficients of r=0.18 (*p* = 0.40) and r=0.58 (*p* = 0.004) respectively. As a result, the frequency domain response of A-HER at 6 Hz, with stimulation durations of 20s, demonstrated higher reliability than the frequency domain response of A-SSR at 6 Hz with the same stimulation durations, and the time domain response of MMN at 0.5 Hz with stimulation durations of 80 seconds.

### Experiment 2

#### A-HER compared with A-ALR and A-SSR

In contrast to A-SSR, A-HER utilizes two types of stimuli, termed as standard (***S***) and deviant (***D***), in its paradigm. The purpose of Experiment 2 is to discern whether the response of A-HER is evoked by the discrepancy among stimuli or the inherent uncertainty in the stimulus sequence. To achieve this, a deterministic sequence known as Auditory Alternative Response (A-ALR), characterized by maximum discrepancy, is utilized for comparison against A-HER (exhibiting maximum uncertainty) and A-SSR (featuring zero discrepancy and zero uncertainty). The stimulation frequency has been fixed at 6 Hz.

The time domain responses of A-HER are compared with both A-ALR and A-SSR in Fig. 4A. The averaged peak-to-peak amplitudes of A-HER, A-ALR, and A-SSR are 3.91 *μ*V, 3.11 *μ*V, and 2.09 *μ*V respectively. The amplitude of A-HER is nearly twice as large as that of A-SSR, while the amplitude of A-ALR falls between A-HER and A-SSR. The paired-sample *t*-test results indicate a significantly larger amplitude for A-HER than A-ALR (*p* = 2.01 × 10^−5^), and for A-ALR than A-SSR (*p* = 2.85 × 10^−4^).

**Figure 4.**
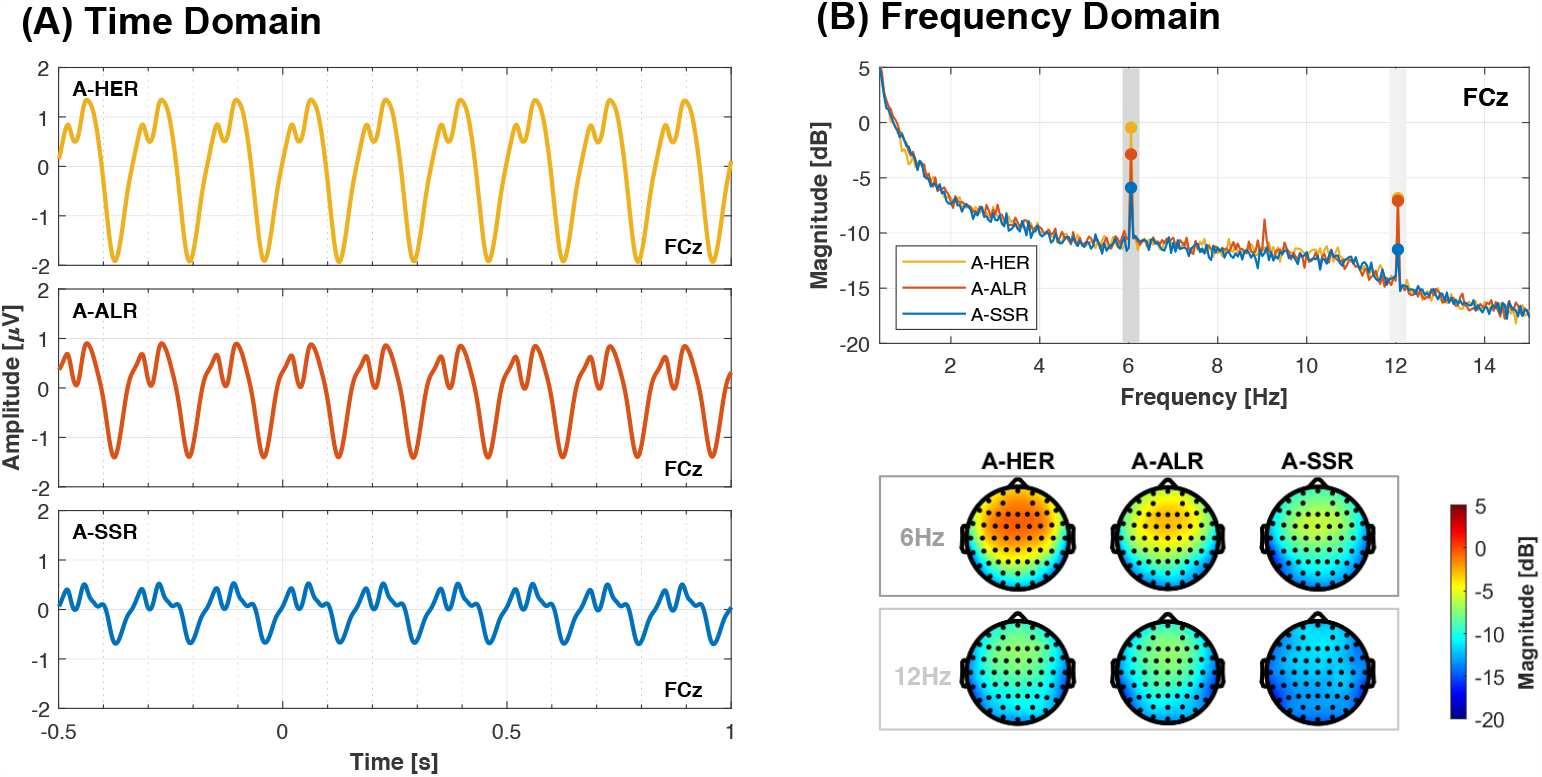
The response of A-HER, A-ALR, and A-SSR with the stimulation frequency of 6 Hz from both (A) time domain and (B) frequency domain in Experiment 2.

The frequency domain analysis, as depicted in Fig. 4B, corroborates these findings. At the fundamental frequency of 6 Hz, the magnitude of A-HER is significantly larger than A-ALR (*p* = 4.94 × 10^−4^), and A-ALR is significantly larger than A-SSR (*p* = 4.54 × 10^−4^). For the harmonic frequency at 12 Hz, there is no significant difference between A-HER and A-ALR (*p* = 0.91). However, both A-HER and A-ALR are significantly larger than A-SSR with corresponding p-values of *p* = 1.17×10^−4^ and *p* = 2.39×10^−6^, respectively. No observable response is detected at 3 Hz for A-ALR. The topographies in Fig. 4B indicate that the responses of A-HER, A-ALR, and A-SSR are primarily concentrated on the channel FCz at both 6 Hz and 12 Hz.

#### Traveling wave analysis

The theta band responses across all channels were portrayed in Fig. 5A. A-SSR manifests a distinct pattern emerging from the occipital lobe and advancing towards the frontal lobe subsequent to auditory stimulation. The topographical map details reveal that at channel FCz in the frontal lobe region (highlighted in red), there is a trough of -0.27*μV* at 129 ms and peaks at 0.22*μV* at 217 ms. Conversely, at channel Oz in the occipital lobe region (depicted in blue), there is a trough of -0.09*μV* at 78 ms and peaks at 0.12*μV* at 180 ms.

**Figure 5.**
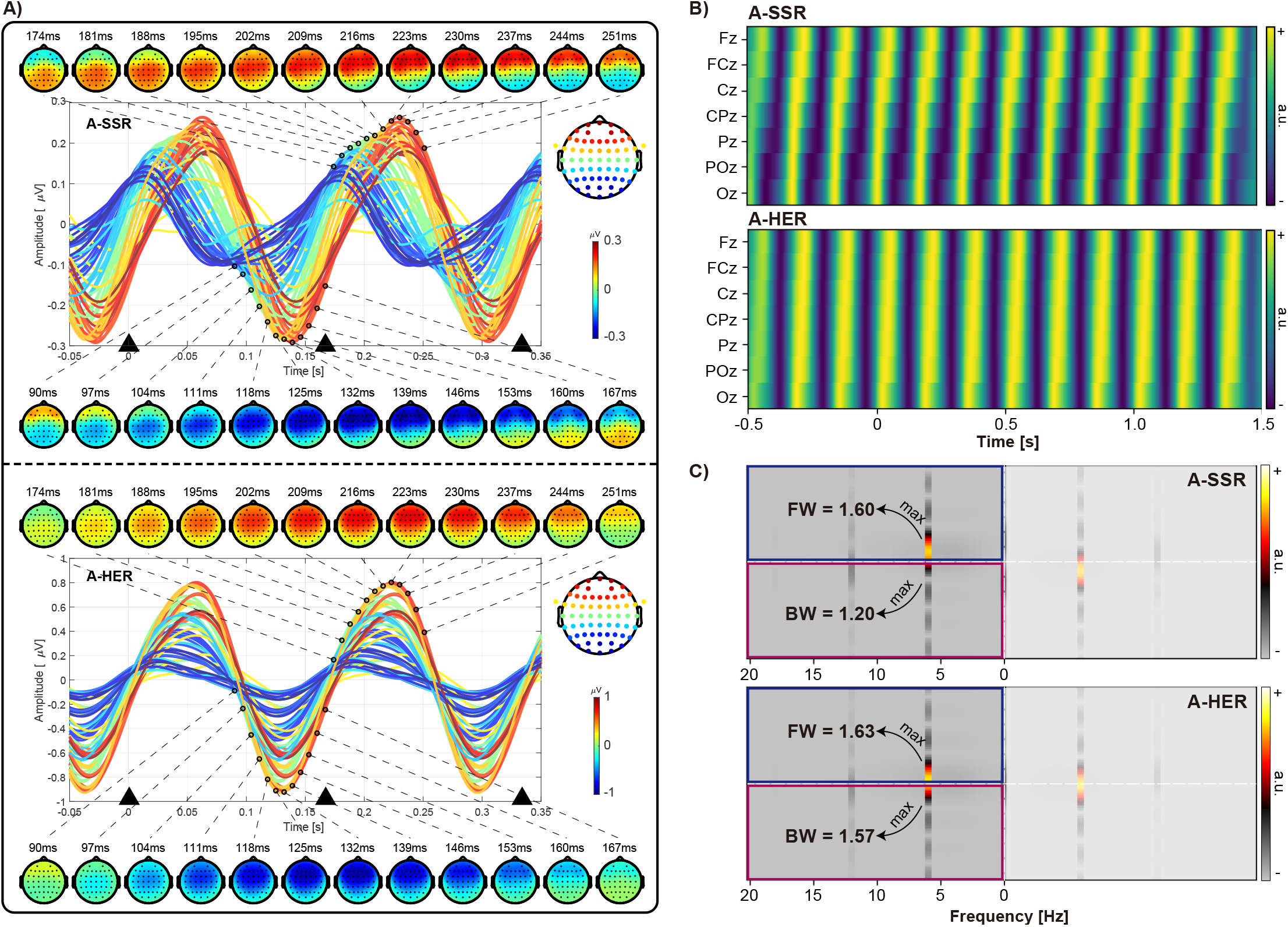
Traveling wave analysis for the theta band in A-SSR and A-HER with the simulation frequency of 6Hz in Experiment 2. (A) The theta band response of A-SSR and A-HER with the 4-8Hz bandpass filter on the EEG signal. The topographies of the peak and trough from 90 - 251 ms are also shown at the top and bottom of the curves. The black triangles “▴” indicate the time points of the auditory stimulation. (B) The normalized 2D-map of A-SSR and A-HER, overlaying data from seven electrodes spanning from the occipital lobe to the frontal lobe (Oz, POz, Pz, CPz, Cz, FCz, Fz). (C) Quantifying traveling wave. Conduculations on the 2D map within the period of -0.5 to 1.5 s. The maximum values in the upper-left and lower-left quadrants respectively indicate the presence of forward (FW) and backward (BW) waves.

In contrast, A-HER lacks these specific traits. Both the peak and trough across all channels occur simultaneously, centralized around the central frontal area near channel FCz. The topographical map delineates that at channel FCz in the frontal lobe region (depicted in red), there is a trough of -0.93*μV* at 128 ms and a peak of 0.78*μV* at 219 ms, while at channel Oz in the occipital lobe region (depicted in blue), there is a trough of -0.16*μV* at 126 ms and peaks at 0.15*μV* at 192 ms. The simultaneous occurrence of peak and trough across all channels is confirmed by visual inspection, showcasing their maximum and minimum values at the central frontal area around channel FCz.

Fig. 5B illustrates the normalized time domain signals of the theta band in both A-SSR and A-HER, arranging electrodes from the occipital lobe to the frontal lobe (Oz, POz, Pz, CPz, Cz, FCz, Fz) to generate a 2D map. Across time, the peaks and troughs of A-SSR display a pronounced tilted angle, while A-HER demonstrates a relatively stable vertical orientation.

To quantify the directional spread of these responses, 2D-FFT was calculated for the 2D map. In Fig. 5C, the maximum value in the upper-left quadrant (depicted in the blue box) represents FW, while the maximum value in the lower-left quadrant (depicted in the blue box) represents BW. The logarithmic ratio of these values provides a measure of the overall wave direction, with positive values primarily indicating FW waves and negative values indicating BW waves. The 2D-FFT analysis depicted in Fig. 5C reveals a pronounced amplification of the signal at frequencies 6Hz and 12Hz, with the maximum values concentrated at the 6Hz frequency. A-SSR demonstrates a markedly higher FW value compared to the BW value, as evidenced by a log-ratio of 0.29 (FW = 1.60, BW = 1.20). In contrast, A-HER exhibits comparable FW and BW amplitudes, reflected in a log-ratio of 0.04 (FW = 1.63, BW = 1.57).

### Experiment 3

Table 1 illustrated the magnitude of the frequency responses in A-HER and A-SSR with different stimulation patterns and frequencies in Experiment 3. The gamma band 44 Hz AM-based stimulation elicited a brain response of -11.82 dB for A-SSR at channel FCz, which was significantly higher than the -17.80 dB for A-HER (*p* = 1.58 × 10^−9^). With the theta band 6 Hz AM-based stimulation, the magnitudes of A-SSR and A-HER were -5.75 dB and -5.60 dB, respectively. The paired-sample *t*-test revealed no significant difference between them (*p* = 0.85). However, when using Tone-based stimulation instead of AM-based stimulation, the response of A-SSR (−5.89 dB) remained the same with no statistically significant difference (*p* = 0.89) at 6 Hz, while A-HER increased to -0.46 dB, which was significantly higher than A-SSR (*p* = 1.52 × 10^−5^). When compared to the conventional A-SSR response of -11.82 dB with 44 Hz AM-based stimulation, the proposed A-HER paradigm could elevate the brain rhythm response to -0.46 dB with 6 Hz Tone-based stimulation (*p* = 8.07 × 10^−8^).

**Table 1.**
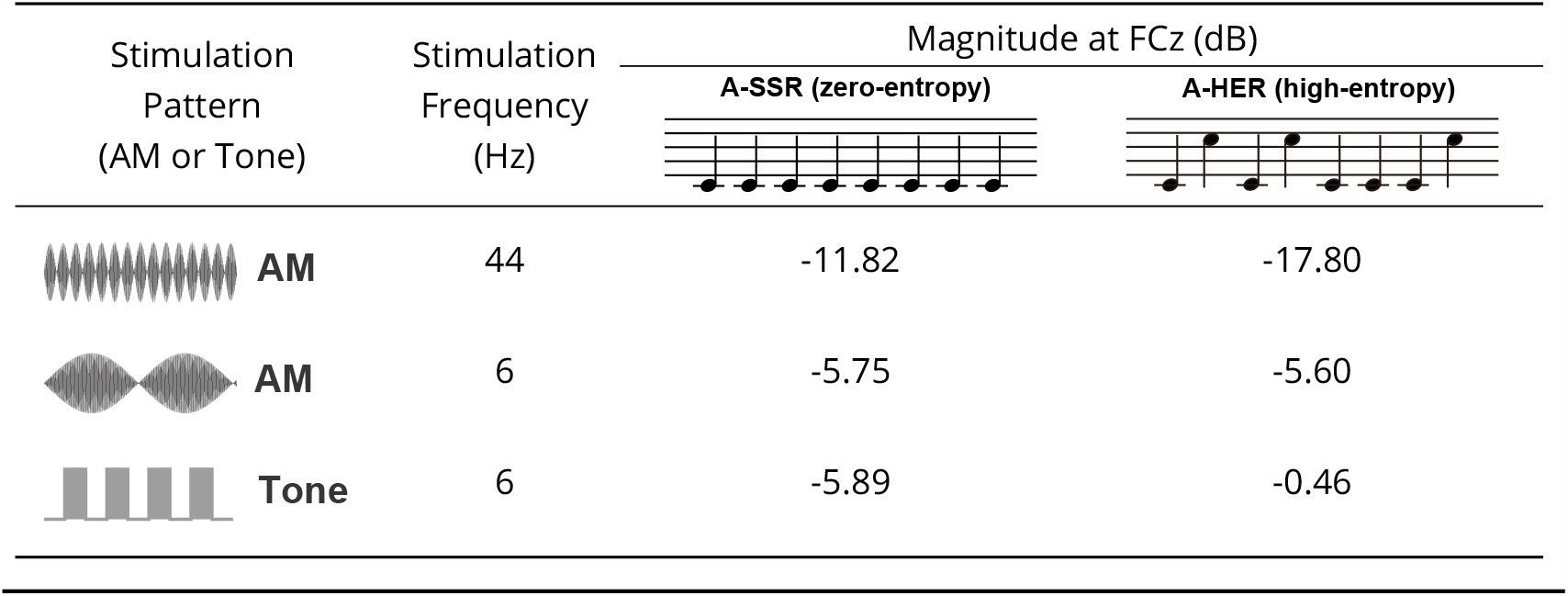
The frequency response of A-SSR and A-HER with different stimulation patterns (AM for amplitude modulated and Tone for pure-tone bursts) and stimulation frequencies (6 Hz, 6 Hz, and 44 Hz) in Experiment 3.

| Stimulation<br>Pattern<br>(AM or Tone) | Stimulation<br>Frequency<br>(Hz) | Magnitude at FCz (dB) |  |
| --- | --- | --- | --- |
|  |  | A-SSR (zero-entropy) | A-HER (high-entropy) |
| **AM** | 44 | -11.82 | -17.80 |
| **AM** | 6 | -5.75 | -5.60 |
| **Tone** | 6 | -5.89 | -0.46 |

## Discussion

### Conclusion

The novel experimental paradigm, Auditory High Entropy Response (A-HER), introduced in this study represents a significant advancement in investigating uncertain information processing within the brain. By prioritizing holistic responses to auditory stimulus sequences with maximized information entropy, A-HER transcends the limitations of conventional paradigms. Our findings reveal that A-HER elicits a substantial frontal theta band response in both the time and frequency domains, markedly enhancing SNR and response reliability compared to traditional zero-entropy A-SSR and low-entropy MMN paradigms. This heightened reliability and SNR position A-HER as a valuable tool for potential applications in neuroengineering, particularly in diagnosing and researching neurological and mental diseases, as well as in advancing brain-computer interfaces. Furthermore, our exploration underscores that the increased magnitude of A-HER response is influenced by both stimulus sequence differences and uncertainties. Notably, while A-HER exhibits a larger amplitude than A-SSR, our research identifies distinct propagation rules and differing principles between evoked and entrained responses. These observations point toward distinct information processing mechanisms for certainty and uncertainty in the brain. In summary, the A-HER paradigm, by surmounting traditional experimental boundaries, provides a pivotal pathway to unraveling how the brain navigates uncertainty in dynamic and unpredictable real-world information scenarios.

### The response of A-HER

The A-SSR has diverse applications in clinical, research, and technological domains, offering insights into auditory function and brain processing mechanisms. While employed for hearing threshold assessment, auditory processing disorders diagnosis, neuroscience studies, and objective hearing screening and BCI applications, A-SSR faces limitations due to its small SNR. This constraint impedes its effectiveness in detecting subtle neural responses amidst background noise, posing challenges in reliable brain activity interpretation, potentially limiting its utility in precision diagnostics and neuroengineering.

With the newly proposed A-HER paradigm, the larger time domain and frequency domain response in frontal lobe with theta band stimulation is the main contribution in this work. Compared to the conventional A-SSR response (44 Hz, AM), the proposed A-HER paradigm (6 Hz, Tone) increased the brain rhythm response from -11.82 dB to -0.46 dB in Experiment 3 (Table 1). The significant 11dB enhancement facilitates quicker detection, leading to notably higher reliability (r = 0.58) within a 20-second stimulus duration when contrasted with both the 20-second A-SSR stimulus and the conventional 80-second MMN test observed in Experiment 1 (Fig. 3). The considerable improvement in SNR observed in A-HER holds promise for its practical application in enhancing neural signal detection and processing. Whether the magnitude can be compared with V-SSR still needs further optimization and validation.

### The evoking of A-HER

#### Evoked responses and neural entrainment

A-SSR relies on amplitude modulation (AM) at 44 Hz, inducing a neural response in the brain that synchronizes with this rhythmic external stimulus. The relatively robust brain response (−11.82 dB), reflected in the measured dB level, indicates the brain’s ability to align and synchronize with this specific regular modulation pattern—a classical manifestation of “neural entrainment”.

Conversely, A-HER, devoid of a regular external rhythmic stimulus, exhibits a comparatively weaker brain response (−17.80 dB, Table 1). Notably, as the stimulation frequency shifts from the gamma band (44 Hz) to the theta band (6 Hz), both A-SSR (−5.60 dB) and A-HER (−5.75 dB) demonstrate similar responses with amplitude modulation. For this reason, we can infer that this phenomenon of neural entrainment occurs mainly in the gamma frequency band.

Moreover, the transition from amplitude modulation (AM) to pure tone burst (Tone) fails to enhance the A-SSR response, while significantly improving A-HER’s response from -5.75 dB to -0.46 dB. This highlights a distinction: A-SSR relies on neural entrainment to a rhythmic pattern, while A-HER’s response is more evoked by individual events within an uncertain sequence. However, this event-driven processing is influenced by the uncertainty of the entire stimulus sequence, indicating a more intricate neural mechanism. This complexity delineates A-HER’s neural response from AEP evoked by isolated stimuli and A-SSR, wherein neural rhythms are synchronized.

#### Uncertainty and discrepancy

The findings from Experiment 2 (Fig. 2) indicate that the amplified amplitude in the A-HER compared to the A-SSR can be linked to two factors: discrepant stimuli and uncertainty. Regarding the A-ALR, it was observed that deterministic discrepant stimuli successfully heightened the amplitude. However, in the case of the A-HER, despite a stimulus difference of less than half compared to the A-ALR, there was a notable enhancement in the stimulus’s amplitude response.

A more in-depth analysis suggests that within the A-HER context, even with a smaller stimulus difference, the amplitude response saw further improvement. This might suggest that the A-HER exhibits heightened sensitivity to both stimulus disparities and uncertainty. Moreover, the role of uncertainty might hold more significance in the A-HER scenario, where even a smaller stimulus difference might lead to a higher cognitive load, consequently amplifying the amplitude response. Nevertheless, further experiments and analysis are essential to validate and elucidate these differences between A-HER and A-ALR.

### The propagation of A-HER

In Experiment 2, discernible differences emerged between A-HER and A-SSR responses, not solely in their magnitudes but also in their propagation patterns.

The first possible explanation comes from the propagation model for travel waves and standing waves (***Gonzalez-Castillo, 2022***). According to the definition, a standing wave can form when two waves of equal amplitude and frequency are traveling in opposite directions; and a traveling wave, as the name implies, is a wave that is moving. Hence, the traveling wave-like results in A-SSR imply that the response of A-SSR has only one source, which may mainly come from STG as the primary auditory area (***Halder et al., 2019***). But why the propagation direction is from posterior to frontal is still hard to explain. While the standing traveling wave-like results in A-HER imply that the response of A-HER may be composed of the propagation of signals from at least two brain sources. Inspired by MMN (***Doeller et al., 2003; Garrido et al., 2008; Tse et al., 2006***), the two possible brain sources would be Heschl’s gyrus located in the posterior portion of the STG as the primary auditory area, and the inferior frontal gyrus (IFG), which is involved in establishing a prediction model for detecting unexpected changes (***Lui et al., 2021***). One counterargument against this propagation model is that if A-SSR adheres to this propagation law, a propagation time of around 40ms from a peak at Oz (180ms) to a peak at FCz (217ms) appears relatively sluggish.

An alternative interpretation proposes that neither A-SSR nor A-HER distinctly exhibit spatial propagation in their theta rhythm responses. Instead, both concurrently involve components from the occipital and frontal lobes. The variation lies in A-HER’s stronger frontal lobe amplitude compared to A-SSR. Both display similar occipital lobe responses without notable divergence. Evidence from channel FCz supports this, with A-HER showing similar peak and trough latencies as A-SSR, albeit with significantly higher amplitudes. At channel Oz, peak amplitudes and latencies in both A-HER and A-SSR align, but the frontal lobe’s influence notably affects trough amplitudes and latencies. A possible reason for channel Oz’s unaffected peak could be its latency aligning near channel FCz’s zero-crossing point, experiencing comparatively less influence. Nonetheless, this interpretation struggles to explain why, as seen in Fig. 5B, A-SSR’s waveform continues propagating forward post-peak at channel FCz.

It’s possible that both explanations oversimplify the scenario. Further investigation, possibly using neuroimaging or localization techniques, is essential to validate these hypotheses. Gaining a deeper understanding of how both STG and IFG contribute to generating the A-HER response could provide a more comprehensive insight into the intricate mechanisms involved in auditory processing.

### Limitation

Firstly, in this study, the pitch of stimuli has not been strictly controlled. It should be noticed that in Fig. 3(A), the response of deviant stimuli ***D*** is always higher than the standard stimuli ***S*** with all stimulation frequencies in A-HER. In Experiment 1, the pitch of stimuli ***S*** and ***D*** is 524 Hz and 262 Hz correspondingly. The result could be explained by the global effect of MMN in the experiment. While we did not switch pitches of stimuli in this experiment for strict control. As a result, the local and global impacts of MMN have not been thoroughly examined, which does not alter our results for investigation of A-HER.

Secondly, in the experiment, 6 Hz was used as the representative stimulation frequency in theta band for the experiment design and EEG signal analysis, which is determined by the results of pre-experiments with a small group of subjects. However, the results of Experiment 1 indicated the response of 7 Hz is larger than 6 Hz. But the analysis of the EEG signal is still focused on 6 Hz. But whether the frequency is 6Hz or 7Hz, the response is focused in the theta band and its features remain constant.

### Future work

Firstly, for brain source localization, further analysis should be done on the comparison of A-HER, A-SSR, and MMN. In this work, A-HER is proposed by using a high uncertainty stimulus sequence to induce a larger frequency response than A-SSR. According to our knowledge of MMN, this response is not only caused by the auditory primary sensory cortex, but also reflects higher cognitive functions are most likely involved in the cognitive processing of this stimulus sequence. Hence, further work on brain source localization is in progress, and we hope that the sEEG and MEG results will provide more evidence about the origins of A-HER.

Second, numerous approaches for increasing the entropy of the stimulus sequence should be explored in order to boost the response of A-HER. In fact, this work just demonstrated the feasibility of A-HER. The result in Fig. 8 demonstrates the larger magnitude of A-HER than A-SSR, but the magnitude is still smaller than the well-studied V-SSR. Several methods can be used to further improve the magnitude of A-HER. On one side, considering inter-stimulus variability rather than stimulus intensity as the main cause of MMN, we can further deliver the two types of stimuli in A-HER paradigm with larger differences in the scale of pitch, intensity, and timbre. On the other side, considering more types of stimuli would lead to a higher entropy in Eq. (1). Multiple types of stimuli should be included in the paradigm to improve the uncertainty of the sequence.

Thirdly, the large response magnitude would make A-HER an ideal paradigm for the clinical functional assessment and engineering applications. 1) The large frequency response of A-HER allows us to potentially develop a more valuable tool for auditory function assessment than A-SSR. 2) The processing of uncertain information in A-HER would make the A-HER paradigm suitable for cognitive function assessment and mental illness diagnosis. In the case of schizophrenia, multiple stimuli with short ISI allow A-HER potentially provide more rapid and accurate diagnostic results than MMN. 3) In the field of BCI application, A-HER provides us with a non-visual interaction paradigm. Furthermore, the stimulation of A-SSR could effectively avoid the sensory fatigue caused by V-SSR, because the response of A-HER does not depend on the stimulus intensity itself but on the difference and uncertainty between stimuli. 4) Considering that uncertainty and surprise would evoke musical pleasure (***Cheung et al., 2019***), the proposed A-HER could potentially be used in music therapy (***Chen et al., 2021; Han et al., 2021***).

## Methods and Materials

### Experimental platform

During the experiment, the subjects were seated in a comfortable chair, which is about one meter from the screen and the speaker, and were asked to fixate on a cross at the center of the screen during the experiment with auditory stimulation. We did not ask the subject to pay attention to the auditory stimuli. A speaker (EDIFIER MR4, Edifier Technology Co Ltd. Shenzhen, China) was used to present auditory stimuli. The intensity was set at a comfortable level (75 dB SPL on average) for all subjects as measured by a digital sound level meter (Victor 824, Double King Industrial Holdings Co., Ltd. Shenzhen, China). A 24.5-inch screen (1920×1080) with a 360-Hz refreshing rate (Alienware AW2521H, Miami, USA) was used to present the visual cues.

In Experiment 1 and Experiment 2, Arduino Uno was programmed to release pure-tone bursts (Tone) auditory stimulation for the accurate marker of each burst and avoid the delay on Windows PC. The duration for each Tone stimulus was 40ms. In Experiment 3, amplitude modulated (AM) auditory stimulation and visual stimulation were implemented by Matlab (The MathWorks Inc., Natick, USA), since only frequency domain analysis has performance in Experiment 3 which is not sensitive to the stimulation delay. All three experimental sequences were presented to subjects by Matlab.

For auditory stimulation, 524 Hz (the C one octave higher than the middle C) and 262 Hz (the middle C) sinusoidal signal was used as standard and deviant stimuli. It should be noted that the terms of standard (***S***) and deviant (***D***) stimuli are from the paradigm of MMN. For A-SSR, there are only standard stimuli. For A-HER and A-ALR, both two types of stimuli were presented with the same probability of 50%, but we still termed them as standard (***S***) and deviant (***D***) stimuli.

The continuous EEG signals were recorded using an EEG amplifier (BrainAmp, Brain Products GmbH, Germany) and multichannel EEG caps (64 Channel, Easycap). The signals were recorded at a sampling rate of 1000 Hz by 64 electrodes, placed in the standard 10–20 positions. The electrodes FCz and AFz served as reference and ground, respectively. Before data acquisition, the contact impedance between the EEG electrodes and the scalp was calibrated to be lower than 10 kΩ to ensure the quality of EEG signals during the experiments.

### Signal Processing

#### Signal pre-processing

For EEG pre-processing, the signal was firstly re-reference to TP9/TP10. A 0.1 to 99 Hz 4th-order Butterworth zero-phase filters were applied as the bandpass filtering and a notch filtering was applied for 50 Hz power-line artifact. Then, artifacts produced by eye blinks or eye movements were identified and removed manually by Independent Component Analysis (ICA).

#### Time domain analysis

After signal pre-processing, a 0.4 − 30 Hz 4th-order Butterworth zero-phase filter was applied for time domain analysis. After that, the EEG signals were segmented by the markers of the standard (***S***) and deviant (***D***) stimuli from -500 ms to 1000 ms to obtain the Event-Related Potential (ERP) for each subject. The maximum and minimum values from the interval of 50 ms to 250 ms of the ERP signal were extracted for further statistical analysis. Compared with the conventional ERP analysis, the SOA in this work was shorter than the interval for segmentation. The overlap between the two successive stimuli existed commonly for the time domain analysis. Hence, the pre-stimulus ERP is also informative, which could not be treated as the conventional baseline. Hence, we did not perform baseline correction in the time domain analysis. With the bandpass filter of 0.4 − 30 Hz, the DC component has been removed effectively.

#### Frequency domain analysis

Considering the different durations for each condition from Experiment 1 to Experiment 3, the pre-processed EEG signal with a duration longer than 20 seconds was first segmented into the segment of 20 seconds with no overlap. Then Fast Fourier Transform (FFT) was performed on these EEG segments. The FFT results from the same conditions and the same stimulus frequency from different blocks were averaged. The magnitudes of the foundational frequency and their second harmonics frequency (2 × foundational frequency) were extracted for further statistical analysis.

#### Traveling wave analysis

To quantify the directional propagation of EEG signals, we adopted a wave quantification method ***Alamia and VanRullen (2019***). Given the limited number of electrodes, zero-padding was applied to enhance frequency resolution ***Kim and Im (2018***). This operation not only increased the number of data points but also facilitated spatial interpolation ***Gao et al (2022***), enabling a more detailed examination of spatial relationships between electrodes.

Specifically, based on the time domain analysis results of Experiment 2, we conducted a 4-8Hz bandpass filter on the data corresponding to ASSR and AHER. Subsequently, both sets of data were normalized to mitigate amplitude effects, allowing for clearer observation of spatial signal characteristics. We sequentially stacked the EEG signals from seven electrodes (Oz, POz, Pz, CPz, Cz, FCz, Fz) to generate a 2D-map. To conduct traveling wave analysis, zero-padding was implemented outside the electrode region, expanding the matrix width from 7 to 101. Subsequently, a 2D-FFT transformation was performed on the 2D-map, yielding temporal frequencies along the horizontal axis and spatial frequencies along the vertical axis.

## Acknowledgements

This work was supported by the National Natural Science Foundation of China (No. 62271326), the Shenzhen Science and Technology Program (No. JSGG20210713091811038), Medical-Engineering Interdisciplinary Research Foundation of ShenZhen University.

## Notes

### Competing Interest Statement

The authors have declared no competing interest.

